# AIscEA: Unsupervised Integration of Single-cell Gene Expression and Chromatin Accessibility via Their Biological Consistency

**DOI:** 10.1101/2022.02.17.480279

**Authors:** Elham Jafari, Travis Johnson, Yue Wang, Yunlong Liu, Kun Huang, Yijie Wang

## Abstract

Since the integrative analysis of single-cell gene expression and chromatin accessibility measurements is essential for revealing gene regulation at the single-cell resolution, integrating these two measurements becomes one of the key challenges in computational biology. Because gene expression and chromatin accessibility are measurements from different modalities, no common features can be directly used to guide their integration. Current state-of-the-art methods assume that the number of cell types across the measurements is the same. However, when cell-type heterogeneity exists, they might not generate reliable results. Furthermore, current methods do not have an effective way to select the hyper-parameter under the unsupervised setting. Therefore, applying computational methods to integrate single-cell gene expression and chromatin accessibility measurements remains difficult.

We introduce AIscEA – **A**lignment-based **I**ntegration of single-cell gene **E**xpression and chromatin **A**ccessibility – a computational method that integrates single-cell gene expression and chromatin accessibility measurements using their biological consistency. AIscEA first defines a ranked similarity score to quantify the biological consistency between cell types across measurements. AIscEA then uses the ranked similarity score and a novel permutation test to identify the cell-type alignment across measurements. For the aligned cell types, AIscEA further utilizes graph alignment to align the cells across measurements. We compared AIscEA with the competing methods on several benchmark datasets and demonstrated that AIscEA is more robust to hyper-parameters and can better handle the cell-type heterogeneity problem. Furthermore, we demonstrate that AIscEA significantly outperforms the state-of-the-art methods when integrating real-world SNARE-seq and scMultiome-seq datasets in terms of integration accuracy.

## 1 Introduction

Advances in single-cell high-throughput technologies have enabled us to profile gene expression and chromatin accessibility at the single-cell resolution [1–11]. Integration of the single-cell gene expression and chromatin accessibility measurements shed light on revealing gene regulation for specific cells [12–16]. However, the heterogeneity among single cells presents challenges for such integration [15]. Single-cell gene expression and single-cell chromatin accessibility measure the cells at the transcriptomic and epigenomic layers, respectively. When both measurements are profiled independently, it is difficult to identify the cell-type or cell-cell correspondences across the measurements since they exist in heterogeneous cellular modalities and lack any shared features to integrate them. Single-cell dual-omics sequencing technologies [17, 18] have been developed to tackle this problem by simultaneously profiling gene expression and chromatin accessibility for the same cells. However, most available single-cell gene expression and chromatin accessibility datasets are still profiled independently. Therefore, a reliable computational method is needed to integrate these two kinds of single-cell measurements from different modalities.

Several unsupervised integrative methods have been developed to integrate the single-cell gene expression and chromatin accessibility measurements [19–27]. CoupleNMF [19] unitizes the non-negative matrix factorization framework to integrate the single-cell gene expression and chromatin accessibility measurements at the cell type level. Other state-of-the art methods focus on the integration at the cell-cell level. They assume that single-cell gene expression and chromatin accessibility measurements share similar low-dimensional manifolds and apply different computational methods to align the corresponding manifolds. MMD-MA aligns the manifold of single-cell gene expression profile and the manifold of single-cell chromatin accessibility profile by minimizing the maximum mean discrepancy between them [23]. UnionCom relies on the generalized unsupervised manifold alignment and uses local and global properties of the cells to align the single-cell gene expression and chromatin accessibility measurements [24]. SCOT applies the Gromov-Wasserstein-based optimal transport to align the manifolds [25], but Pamona unitizes the partial Gromov-Wasserstein optimal transport to align the manifolds [26].

However, all current methods mentioned above suffer from two major problems. First, they are incapable of handling the cell-type heterogeneity problem. Specifically, when the cell types in the single-cell gene expression profile differ from those in the single-cell chromatin accessibility profile, they may generate poor alignment. CoupleNMF [19] would fail because it requires the two datasets have the same number of cell types. Other methods assume that single-cell gene expression and chromatin accessibility share a similar manifold, which might not hold when the cell types across datasets are different. MMD-MA [23], Union-Com [24], and SCOT [25] methods that rely on such assumptions would enforce the alignment between two different manifolds, which would lead to incorrect integration. Pamona [26] attempts to resolve the celltype heterogeneity predicament by estimating the number of the common cells across diverse measurements. However, the performance of the proposed estimation has not been comprehensively tested [26]. Second, all current methods’ performance highly relies on hyper-parameter selection. Under the unsupervised setting, it is very challenging to find the optimal hyper-parameter for different datasets.

To overcome these limitations, we present AIscEA – **A**lignment-based **I**ntegration of single-cell gene **E**xpression and chromatin **A**ccessibility – an unsupervised computational method that explicitly uses biological consistency between gene expression and chromatin accessibility to guide the across-modality integration. First, AIscEA defines a rank-based similarity score to quantify the biological consistency between cell types in gene expression and chromatin accessibility profiles. Then, based on the rank-based similarity and a novel designed permutation test, AIscEA identifies the domain-specific cell types and then finds corresponding cell types shared across single-cell gene expression and chromatin accessibility profiles. Furthermore, for these corresponding cell types across modalities, AIscEA applies a graph alignment method to elucidate the cell-cell correspondence [28].

We first validated the performance of AIscEA using SNARE-seq Human cell line mixtures data [17], which jointly captured accessible chromatin regions and gene expression profiles within the same cells, and therefore it provides cell-cell correspondence for benchmarking. The benchmarking results demonstrate that AIscEA can resolve the cell-type heterogeneity problem and is more robust to hyper-parameters. Furthermore, we show that our AIscEA outperforms CoupleNMF [19] in terms of cell-type alignment. In addition, we compared the performance of our method with state-of-the-art cell-cell integration methods MMD-MA [23], UnionCom [24], SCOT [25], and Pamona [26] on real-world single-cell gene expression and chromatin accessibility profiles. We applied them to integrate SNARE-seq profilings of neonatal mouse cerebral cortex [17], adult mouse cerebral cortex [17], and two scMultiome-seq PBMC datasets from the healthy donors. We demonstrate that AIscEA significantly outperforms other methods in terms of the average FOSCTTM score [23], demonstrating its outperformance in identifying the cell-cell correspondence.

## 2 Methods

### 2.1 Method Overview

AIscEA is an alignment-based method that can identify the cell-type and cell-cell correspondence between single-cell gene expression and chromatin accessibility measurements profiled from the same tissue. In contrast to the current state-of-the-art methods [23–26], AIscEA does not rely on the assumption of similarity between the manifolds of the entire single-cell gene expression and chromatin accessibility measurements. However, AIscEA relies on biological consistency, which is the fact that the promoter regions of over-expressed genes should be significantly accessible to guide the alignment between cell types and also between cells across the measurements [29–33]. AIscEA quantifies such biological consistency using a rank-based similarity score and further unitizes the similarity score to direct the cell-type and cell-cell alignments. As shown in Fig. 1, AIscEA consists of three steps: (i) cell-type identification, (ii) cell-type alignment, and (iii) cell-cell alignment. In the following, we will elaborate on the details of each step.

**Fig. 1:**
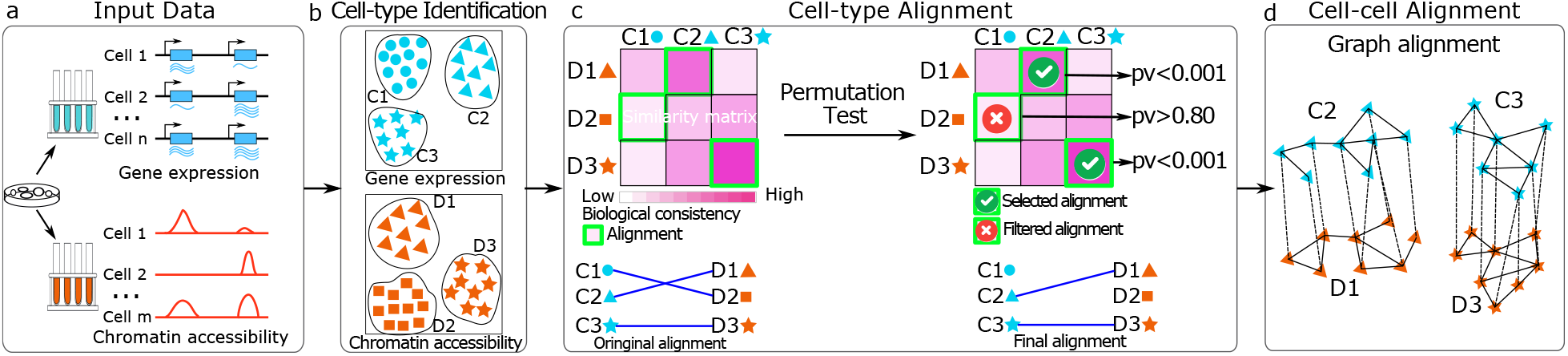
Overview of AIscEA. (a) Input datasets of single-cell RNA-seq and single-cell ATAC-seq measurements. (b) Clustering and cell-type identification. (c) Cell-type alignment using biological consistency and calculating the p-values by a novel permutation test. (d) AIscEA finds cell-cell alignment using a graph alignment method for each pair of mapped clusters.

### 2.2 Cell-type Identification

We first identify cell types within single-cell gene expression and chromatin accessibility measurements via commonly used clustering methods. For single-cell gene expression, we use the classical graph-based clustering method [34–37] to identify *n* cell-types in 𝒞 = {*C*_1_, *C*_2_, …*C*_*n*_}. For single-cell chromatin accessibility measurement, we first use cisTopic [38] to extract regulatory topics and then use the extracted features to cluster cells into *m* cell-types 𝒟 = {*D*_1_, *D*_2_, …*D*_*m*_}. In practice, we use the Leiden clustering as the default clustering method to identify cell types in both single-cell gene expression and chromatin accessibility measurements.

Furthermore, for each cell-type *C*_*i*_, ∀*i* identified in the single-cell gene expression measurement, AIscEA identifies the set of differential over-expressed genes 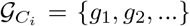. AIscEA then ranks these genes by their expression log 2 fold changes with respect to their expression in the rest of the cells in descending order. We further use a function 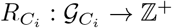 to retrieve the ranking of a gene in 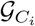. Similarly, for each cell-type *D*_*j*_, ∀*j* identified in the single-cell chromatin accessibility measurement, AIscEA identifies the significantly accessible locations using the predictive distribution calculated by cisTopic [38]. Next, we identify the overlap between these significantly accessible locations and the promoter regions of the expressed genes. AIscEA uses 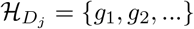 to present the set of genes, whose promoter regions overlap with the significantly accessible locations in cell-type *D*_*j*_.

### 2.3 Cell-type Alignment

After cell-type identification, as explained in section 2.3, *n* and *m* cell types are obtained in the single-cell gene expression and chromatin accessibility measurements, respectively. Although both measurements are profiled from the same tissue, due to cellular heterogeneity, in general, the number of cell types *m* and *n* may differ. Furthermore, the correspondence is unknown between *n* cell-types in the single-cell gene expression measurements and *m* cell-types in the single-cell chromatin accessibility measurements.

We propose to use the biological consistency between gene expression and chromatin accessibility to find the alignment of cell types across different modalities (as shown in Fig. 1). Furthermore, AIscEA adopts a novel permutation test to find statistically significant biological consistency between the aligned cell types. We will keep the aligned cell types that are statistically significant and filter out the rest, as shown in Fig. 1c.

#### Cell-type alignment by biological consistency

The biological consistency AIscEA anchored on is the fact that the promoter regions of over-expressed genes should be significantly accessible [29–33]. AIscEA defines a ranked similarity score *S* to quantify such biological consistency between cell types. Mathematically, the ranked similarity score *S*(*C*_*i*_, *D*_*j*_) between cell-type *C*_*i*_ in single-cell gene expression and cell-type *D*_*j*_ in single-cell chromatin accessibility data can be computed by:

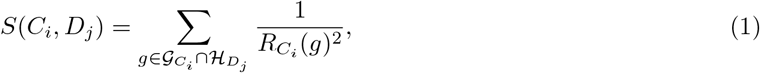

where 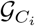 is the set of differential over-expressed genes in cell-type *C*_*i*_ identified in the single-cell gene expression measurements. 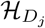 is the set of genes whose promoter regions are significantly accessible in cell-type *D*_*j*_ in single-cell chromatin accessibility measurements. 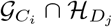 extracts all differentially over-expressed genes whose promoter regions are significantly accessible. 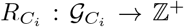 is the function that takes a gene and returns the ranking of the gene in terms of its expression log 2 fold change. The larger the log2 fold change of a gene expression is, the higher rank it has (the rank of the top gene is 1). Based on the definition of *S*(*C*_*i*_, *D*_*j*_) in (1), we know that *S*(*C*_*i*_, *D*_*j*_) is large when 1) 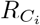(*g*) is small, meaning the top ranking genes’ promoter regions should be significantly accessible; 2) 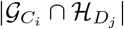 is large, meaning most of the highly over-expressed genes should have significantly accessible promoter regions. Fig. 2(a-c) illustrates a toy example of how *S*(*C*_*i*_, *D*_*j*_) is computed.

**Fig. 2:**
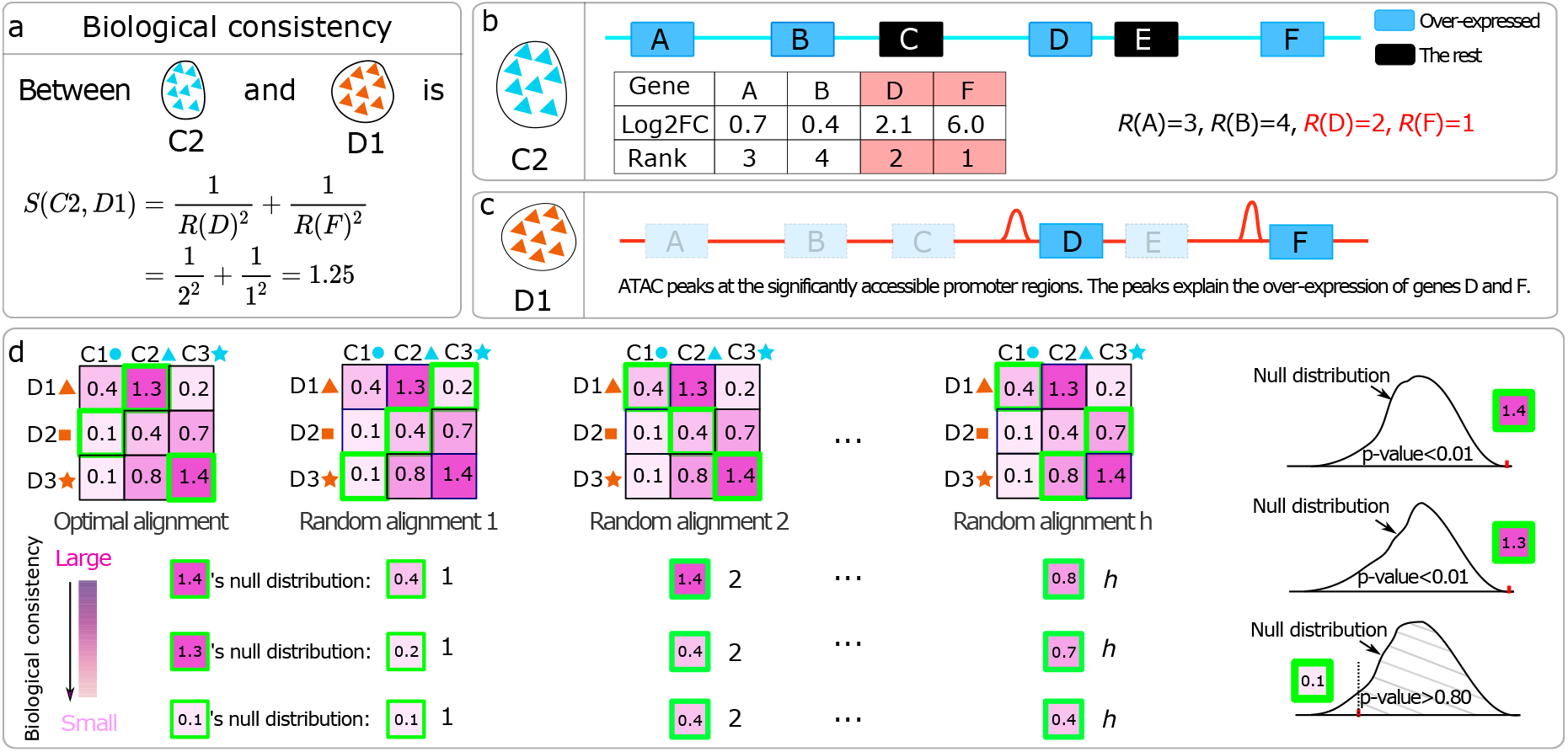
(a) An illustration of computing the biological consistency between (b) a cell type in gene expression, (c) and a cell-type in chromatin accessibility measurement. (d) An explanation of the proposed permutation test to calculate p-values for aligned cell types.

From (1), we compute the biological consistency between *n* cell-types in 𝒞 = {*C*_1_, *C*_2_, …*C*_*n*_} and *m* cell-types in 𝒟 = {*D*_1_, *D*_2_, …*D*_*m*_}. Without loss of generality, we assume that *n* ≤ *m* (if *n* ≥ *m*, we can add dummy cell-types in 𝒟 to make *n* = *m*). Then the cell-type alignment across measurements can be obtained by maximizing the biological consistency between aligned cell types across measurements, which can be formulated as a linear assignment problem:

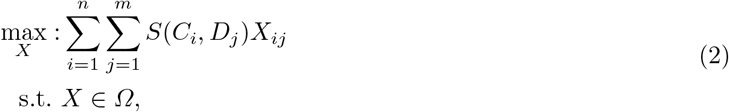

where *X* is a binary assignment matrix, where *X*_*ij*_ = 1 denotes that cell-type *C*_*i*_ corresponds to cell-type *D*_*j*_. The constraint set *Ω* ={*X* ∈{0, 1} ^*n×m*^ : *X***1**_*m*_ = **1**_*n*_, *X*^T^**1**_*n*_≤**1**_*m*_} enforces each cell-type in 𝒞 is assigned to one and only one cell-type in 𝒟. The linear assignment problem can be efficiently solved by the Hungarian algorithm [39].

#### Resolving the cell-type heterogeneity problem via a novel permutation test

The set of cell types 𝒞 in single-cell gene expression could be different from the set of cell types 𝒟 in the single-cell chromatin accessibility data, which results in the cell-type heterogeneity problem. To elucidate the cell-type heterogeneity across measurements, we develop a novel permutation test to distinguish statistically significant corresponding cell types across modalities and find the unique cell types within the measurements.

Before introducing the permutation test, let us first introduce some notations. Given an assignment matrix *Z* ∈ *Ω*, we can obtain the corresponding ranked similarity scores for each alignment and collect them in the set 𝒮_*Z*_ = {*S*(*C*_*i*_, *D*_*j*_)|*Z*_*ij*_ = 1, ∀*i, j*}. We further sort the ranked similarity scores in 𝒮_*Z*_ in descending order and define a function 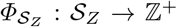 that returns the ranking of a similarity score in 𝒮_*Z*_. We also define 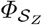 ’s inverse function 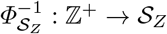 that applies to a given ranking and returns the corresponding similarity score.

The null hypothesis of our novel permutation test is that the ranked similarity score between the aligned cell-types found by (2) are greater or equal to the ranked similarity scores of the randomly aligned cell types. After solving (2), we obtain an optimal assignment *X*\* and the corresponding similarity scores 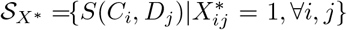. For a specific alignment 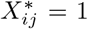, we can get the corresponding similarity score *S*(*C*_*i*_, *D*_*j*_) and its ranking among all alignments by 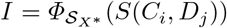. We then generate *k* random cell-type alignments by uniformly sampling *h* = 1, 000 assignment matrices *Z*_1_, …, *Z*_*h*_ ∈ *Ω* from *Ω*. The null distribution of the *I*th ranking similarity score can be estimated by 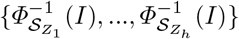 (where 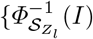 is the *I*th ranked similarity score in the random alignment *Z*_*L*_). By comparing *S*(*C*_*i*_, *D*_*j*_) with 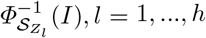, we can calculate the *p*-value by 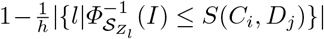, where |·| is the cardinality of a set. For the alignment 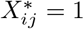 whose corresponding *p*-value is significant (≤ 0.01), we consider it a true alignment. For the alignment 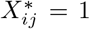 whose corresponding *p*-value is not significant (*p*-value¿0.01), we consider the corresponding cell-types in this alignment may be considered unique cell-types within their measurement. Fig. 2 (d) illustrates how the permutation is calculated.

#### Hyper-parameters selection scheme for the cell-type alignment

Due to the heterogeneity between the single-cell gene expression and chromatin accessibility measurements, the number of cell types *n, m* found in both measurements typically differs (*n* ≠ *m*). The selection of *n* and *m* would influence the performance of the cell-type alignment in AIscEA. Currently, under the unsupervised setting, there is no effective way to select *n* and *m*. To fill the gap, we propose a heuristic approach to select them. AIscEA applies Leiden clustering [35] to identify cell types using the resolution parameter. Therefore, we propose an effective scheme to select the resolution parameter rather than the number of cell types as following.

Our heuristic approach sets the range for the resolution parameter *r*_*e*_ for single-cell gene expression measurement *r*_*e*_ ∈ {0.1, 0.15, 0.2, …, 1.5} and the resolution parameter *r*_*c*_ for single-cell chromatin accessibility measurement *r*_*c*_ ∈ {0.1, 0.15, 0.2, …, 1.5.} Then we screen different combinations of *r*_*e*_ and *r*_*c*_ to compute the *alignment ratio* defined as 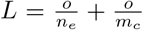, where *n*_*e*_ is the number of identified cell types in gene expression measurement when the resolution parameter is set to *r*_*e*_, *m*_*c*_ is the number of identified cell types in chromatin accessibility measurement when the resolution parameter is set to *r*_*c*_, and *o* is the number of aligned cell types between *n*_*e*_ and *m*_*c*_ identified by the cell-type alignment method in AIscEA (as explained in Section 2.3). In the end, after screening all resolution parameters, we select the ones yielding the largest *alignment ratio L*.

In the experiment Section 4.1, we empirically show that the proposed heuristic approach can select *r*_*e*_ and *r*_*c*_ that result in descent cell-type alignment results for all datasets in a completely unsupervised manner.

### 2.4 Cell-cell Alignment

Once we identify cell-types *C*_*i*_ and *D*_*j*_ are aligned together, we can further find the cell-cell correspondence between the cells in *C*_*i*_ and *D*_*j*_. AIscEA assumes that *C*_*i*_ and *D*_*j*_ consist of a set of cells 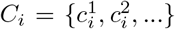 and 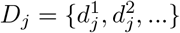, respectively. Since the gene expression of cells in *C*_*i*_ and the chromatin accessibility of cells in *D*_*j*_ are considered as different measurements for cells of the same kind, we confidently assume that their low-dimensional manifold is similar. Hence, a graph alignment method is employed to find the cell-cell correspondence [28].

AIscEA constructs a symmetric *k*-nearest neighbor graph *G*_1_ = (*V*_1_, *E*_1_) to present the manifold of the cells in the cluster *C*_*i*_, where vertices in 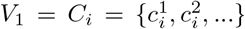 are cells in *C*_*i*_. Similarly, we construct a symmetric *k*-nearest neighbor graph *G*_2_ = (*V*_2_, *E*_2_) to present the manifold of the cells in *D*_*j*_, where vertices in 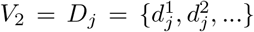 are cells in *D*_*j*_ (details in the Supplementary Materials Section B). Therefore, in the following, we can safely assume |*V*_1_| = |*V*_2_| = *N*. If |*V*_1_| ≠ |*V*_2_|, we can add dummy node to make them equal as done in [28]. The manifold matching between cells in *C*_*i*_ and the cells in *D*_*j*_ can be achieved by the graph alignment between *G*_1_ and *G*_2_. Mathematically, the graph alignment step in AIscEA can be formulated [28] as:

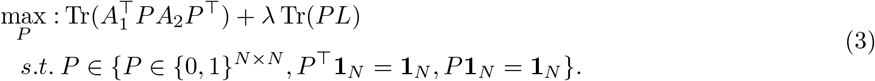

*A*_1_ and *A*_2_ are the adjacency matrices for *G*_1_ and *G*_2_, respectively. *P* is constrained to be a permutation matrix that enforces one-to-one mapping between cells in *G*_1_ and *G*_2_. *L* is the similarity matrix between cells in *V*_1_ and *V*_2_ and *L*_*kl*_ estimates the biological consistency between cells 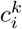 and 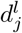. *L*_*kl*_ can be computed by a ranked similarity score, which is similar to (1) (details in the Supplementary Materials). In the objective function in (3), the first term 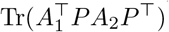 computes the number of overlapping edges between *G*_1_ and *G*_2_ (more overlapping edges imply that the manifolds represented by *G*_1_ and *G*_2_ are similar) and the second term computes total similarity between the aligned cells. *λ* is a hyper-parameter that balances the trade-off in the objective function (3). The optimization in (3) finds a one-to-one cell-cell alignment such that the number of overlapping edges between *G*_1_ as well as *G*_2_ and the total similarity between the aligned cells are maximized simultaneously. We propose applying the Frank-Wolfe [40] algorithm and the path-relinking technique to solve (3) (details in Supplementary Materials Section B).

#### Hyper-parameters for the cell-cell alignment

There are two hyper-parameters that needs to be selected for the cell-cell alignment used in AIscEA: *k* (the number of nearest neighbors when constructing the symmetric *k*-nearest neighbor graph) and *λ* (the regularizer in Eq. 3). In the experiment section 4.1, we show that the cell-cell alignment in AIscEA is robust to the selection of *k* and *λ*. Therefore, we set *k* and *λ* to default values in practice.

## 3 Experimental Setup

### 3.1 Competing methods

AIscEA can identify cell-type alignment between scRNA-seq and scATAC-seq datasets, therefore, we compare AIscEA’s performance on cell-type alignment with CoupleNMF [19], which is the state-of-the-art cell-type alignment method. In addition, AIscEA is able to find cell-cell alignment, therefore, we compare AIscEA with the current state-of-the-art cell-cell alignment methods MMD-MA [23], UnionCom [24], SCOT [25], and Pamona [26].

### 3.2 Data

*SNARE-seq Human* [17] is a joint profiling of accessible chromatin and RNA of the mixture of human cell lines BJ, H1, K562, and GM12878. We use *SNARE-seq Human* to benchmark the competing methods because it provides the ground truth for both cell-type alignment and cell-cell alignment. Moreover, to evaluate different methods on handling the cell-type heterogeneity problem, we generate *SNARE-seq Human_Heterogeneity* data by manually removing cells of BJ cell type from the scRNA-seq data in *SNARE-seq Human*. Furthermore, To compare different methods’ robustness of hyper-parameter selection, we generate *SNARE-seq Human_R5%* and *SNARE-seq Human_R10%*, where 5% of cells and 10% cells are randomly removed from the original *SNARE-seq Human*. Furthermore, we generate *SNARE-seq Human_Heterogeneity_R5%* and *SNARE-seq Human_Heterogeneity _R10%*, where 5% of cells and 10% cells were randomly removed from the *SNARE-seq Human_Heterogeneity* data.

Additionally, we compare all the competing methods on real-world datasets. We first benchmark our method against all competing methods on two SNARE-seq real-world datasets: *SNARE-seq Mouse 5k* (SNARE-seq of neonatal mouse cerebral cortex that contains 5k cells) and *SNARE-seq Mouse 10k* (SNARE-seq of adult mouse cerebral cortex that has 10k cells). Then we apply all competing methods on two sc-Multiome datasets [41, 42]: *scMultiome PBMC* 3*k* (scMultiome-seq PBMC of a healthy donor with 3*k* cells) and *scMultiome PBMC* 12*k* (scMultiome-seq PBMC of a healthy donor with 12*k* cells). All these datasets provide cell-cell correspondence information, which is used to evaluate the competing methods. More details of these datasets can be found in Supplementary Materials Section C.1.

### 3.3 Metrics

We first introduce the metric we use for evaluating the cell-type alignment. When two cell types are aligned between scRNA-seq and scATAC-seq, we expect the cells in scRNA-seq cell type to appear in the aligned scATAC-seq cell type (for the existing cells). In other words, the two aligned cell types are expected to have a larger number of overlapping cells. Therefore, we use the overlap coefficient to measure the overlap between aligned cell types. Specifically, if cell-type *C*_*i*_ is aligned to cell-type *D*_*j*_, the overlap coefficient between cells in *C*_*i*_ and cells in *D*_*j*_ can be computed as:

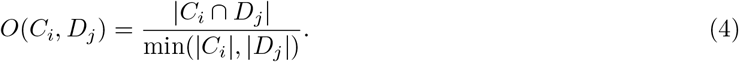

Furthermore, we can compute the total number of the overlapped cells over all aligned cell types as:

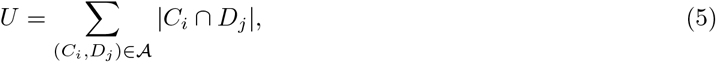

where 𝒜 is the collection of all aligned cell types. Another metric we use to evaluate the cell-type alignment is the average Silhouette score per cluster to measure cluster cohesion. We expect the cells in the same cell types to be similar to other cells in their own cell type, but different from other cell types.

For evaluating the cell-cell alignment, we use the average FOSCTTM score [23], which has been widely used for evaluating single-cell multi-omics integration methods [23, 25, 26]. FOSCTTM stands for “fraction of samples closer than the true match”, therefore, the lower the better. The details of how FOSCTTM is computed are elaborated in Supplementary Materials Section D.1. Another metric is the cell coverage, which is the number of cells that have been found correspondence across the scRNA-seq and scATAC-seq datasets.

### 3.4 Hyper-parameter selection

We select the hyper-parameter for AIscEA using the approaches described in Section 2.3 and Section 2.4. For CoupleNMF [19], we set the number of cell-types based on the ground truth and for the rest of the hyper-parameters, we use the suggested hyper-parameters. For MMD-MA [23], UnionCom [24], SCOT [25], and Pamona [26], under the unsupervised setting, they do not have an effective way to find the optimal hyper-parameters. In this paper, we use the following strategy to find the optimal hyper-parameters for them. We performed a grid search to find the optimal hyper-parameters using *SNARE-seq Human* dataset. Then, we use the optimal hyper-parameters found in *SNARE-seq Human* for real-world datasets (more details in Supplementary Materials Section C.5 and C.6).

### 3.5 Computational resource

All experiments are processed on an Intel(R) Core(TM) i7-6850K CPU @ 3.60GHz CPU with 62GB memory and GPU computations on a single GeForce GTX 1080 Ti with VRAM of 11GB. If a method fails to run on a large-scale dataset due to memory shortage, we report a memory error.

## 4 Results

### 4.1 Benchmarking using SNARE-seq human cell line mixtures

SNARE-seq human cell line mixtures provide the ground truth information for validating cell-type alignment and cell-cell alignment. Therefore, we first use it to validate AIscEA’s hyper-parameter selection scheme proposed in Section 2.3 for cell-type alignment. Furthermore, we use it to evaluate all methods’ robustness to the choice of hyper-parameters for the cell-cell alignment. Last but not least, we use it to benchmark the performance of the competing methods on handling the cell-type heterogeneity problem, as in real-world datasets, the number of cell types may differ between two domains.

#### Validation of the hyper-parameter selection scheme for the cell-type alignment in AIscEA

In Section 2.3, we propose an approach to select the resolution hyper-parameters in the Leiden clustering in AIscEA, which determine the number of cell types in scRNA-seq *n*, and the number of cell types in scATAC-seq *m* for the cell-type alignment in AIscEA. This section uses the *SNARE-seq Human* cell line mixtures to demonstrate that our unsupervised parameter selection scheme can select the resolution hyper-parameters that result in promising cell-type alignment.

We applied the proposed scheme in 2.3 to *SNARE-seq Human_R5%* data, *SNARE-seq Human_R10%* data, *Human_Heterogeneity_R5%* data, and *SNARE-seq Human_Heterogeneity_R10%* data (description of these four data can be found in Section 3.2) and show the hyper-parameters screening results in Fig. 5. In the Fig. 5, the size of each dot corresponds to its *alignment ratios* defined in 2.3. Larger size of the dots means the corresponding *alignment ratio* is higher. The color of each dot for each pair of resolution values indicates the number of overlapping cells identified by the cell-type alignment (computed as *U* defined in Section3.3). Darker blue means more number of overlapping cells are identified by the cell-type alignment, which means the performance of the cell-type alignment is more promising. As shown in Fig. 5, the large-size dots always appear in dark blue color, demonstrating that the *alignment ratio* and the performance of the cell-type alignment method is positively correlated. Therefore, we can use the *alignment ratio* to guide the selection of the resolution hyper-parameters used in AIscEA in an unsupervised setting. In addition, we noticed that many dots have the same size and color. Such observation implies that different combinations of resolution parameters may yield equivalently good cell-type alignments. We have the same observation from the screening results for more real-world datasets in the Supplementary Materials Fig. S2.

#### Benchmarking hyper-parameter robustness in the cell-cell alignment

In this section, we compare AIscEA with all competing cell-cell alignment methods in terms of their robustness to the choice of hyper-parameters. Such robustness is of practical importance since the real-world application is completely unsupervised; therefore, we do not have any prior knowledge to guide the hyper-parameter selection. If a method is sensitive to hyper-parameters, its performance is unreliable for real-world applications.

We applied all methods to the *SNARE-seq Human* data. We ran each method over an extensive grid search of suggested hyper-parameters and showed the results for *SNARE-seq Human* in Table 1. Cell coverage for AIscEA and all other methods are exactly 1, 047 cells. The grid search hyper-parameter tuning details are elaborated in Supplementary Materials Section C.2. As shown in the table 1, AIscEA is competitive with SCOT on achieving the smallest FOSCTTM score, which is superior to the rest of methods. However, AIscEA has the smallest standard deviation, implying that AIscEA is more robust to the choice of hyper-parameters. To further confirm the robustness of hyper-parameters for each method, we applied the optimal hyper-parameters found on *SNARE-seq Human* to the datasets that are slightly different from *SNARE-seq Human*.

**Table 1:**
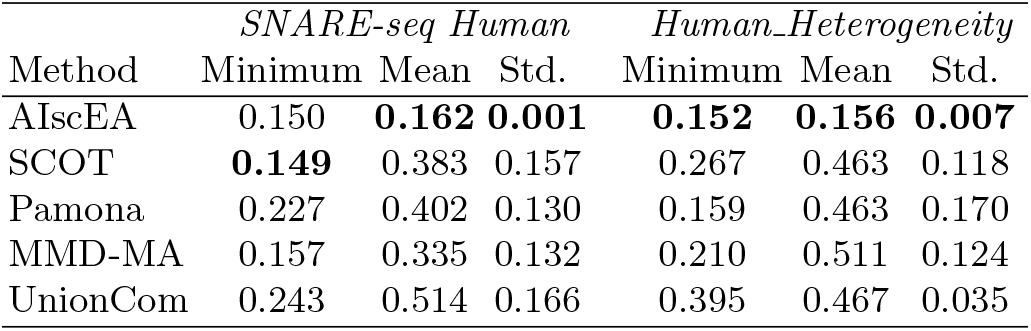
The statistics of average FOSCTTM scores over the grid search of the hyper-parameter for each method using *SNARE-seq Human* and *Human_Heterogeneity* data.

The goal is to check whether the optimal parameters on one dataset would still yield good results on a slightly different dataset.

To evaluate different methods in this scenario, we generated 10 *SNARE-seq Human_R5%* data and 10 *SNARE-seq Human_R10%* datasets (description of the data is in 3.2). Then we applied each method to them using their optimal set of hyper-parameters (in Supplementary Materials Table S1). Cell coverage for AIscEA and competing methods in these experiments is the number of shared cells between two domains for all methods. Fig. 3 c and d exhibit the box plots of the average FOSCTTM scores obtained by each method over 10 *SNARE-seq Human_R5%* data and 10 *SNARE-seq Human_R10%*, respectively. Clearly, our method shows the smallest variance on both figures. We further found that the mean of the average FOSCTTM scores achieved by AIscEA is significantly smaller than the rest of the methods. All these results demonstrate that AIscEA is more robust to the choice of hyper-parameters than all other competing methods.

**Fig. 3:**
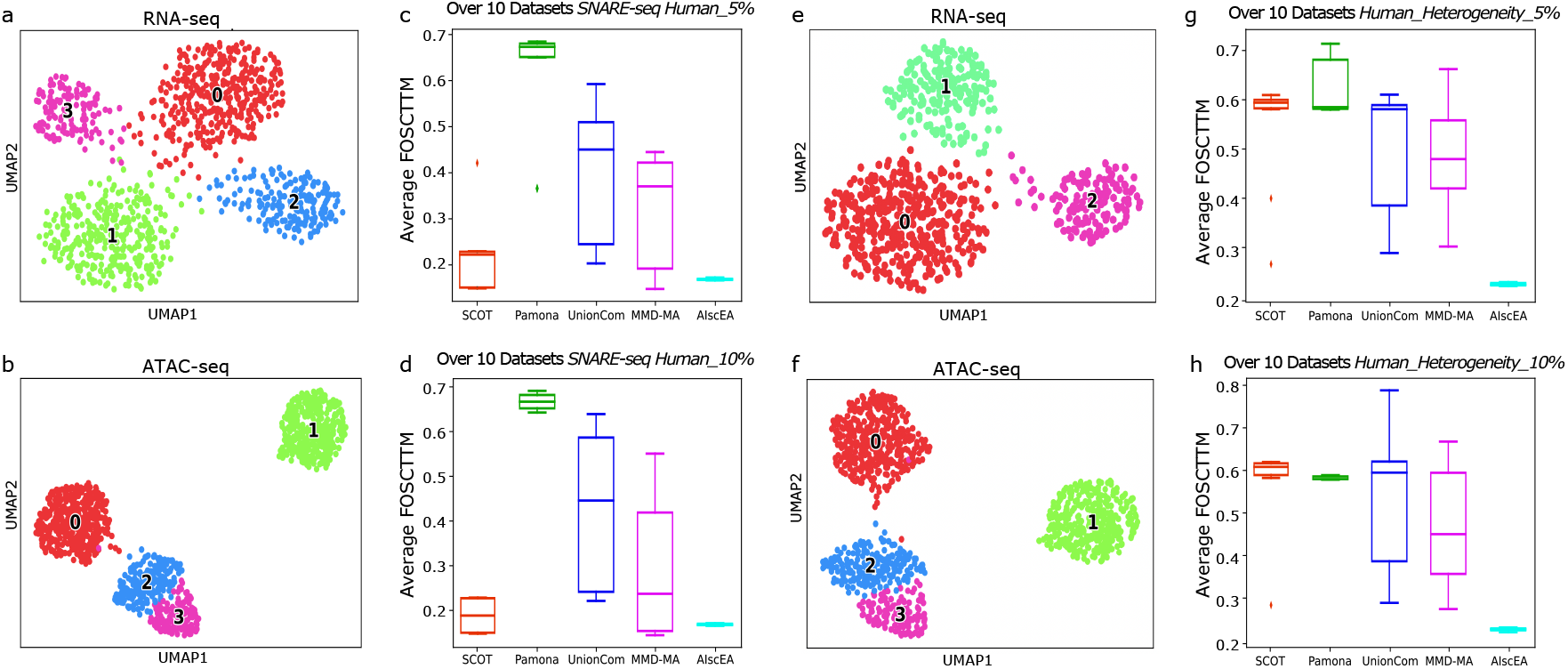
(a) The UMAP of the RNA-seq in *SNARE-seq Human* data. (b) The UMAP of the ATAC-seq in *SNARE-seq Human* data. (c) The box plot of the average FOSCTTM over 10 *SNARE-seq Human_R5%* data. (d) The box plot of the average FOSCTTM over 10 *SNARE-seq Human_R10%* data. (e-f) The UMAPs of SNARE-seq Human_Heterogeneity data that ATAC-seq has one more cell type than RNA-seq. (g) The box plot of the average FOSCTTM over 10 *SNARE-seq Human_R5%* data. (h) The box plot of the average FOSCTTM over 10 *SNARE-seq Human_R10%* data.

#### Benchmarking in solving the cell-type heterogeneity problem

Next, we benchmark all methods on their ability to resolve the cell-type heterogeneity problem. In a real-world application, we may not have any prior knowledge of whether the single-cell RNA-seq measurement and the single-cell ATAC-seq measurement have the same cell types. If we cannot distinguish the cell types that have correspondence and other cell types that have not, the alignment between the two measurements would be misleading.

To simulate the cell-type heterogeneity problem, we generated the *SNARE-seq Human_Heterogeneity* data (description of the data is in 3.2). We first ran each competing method over an extensive grid of suggested set of hyper-parameters and showed the results for *SNARE-seq Human_Heterogeneity* in Table 1. AIscEA can identify the heterogeneous cell type, exclude it, and map the shared cell types between two domains. Cell coverage for AIscEA in this experiment consists of the number of cells in all three shared clusters. As shown, AIscEA achieved the smallest average FOSCTTM score with the smallest standard deviation, indicating AIscEA is the best method to handle the cell-type heterogeneity problem.

Furthermore, we applied each method using its optimal hyper-parameters found on *SNARE-seq Human_Heterogeneity* data (shown in Supplementary Materials Table S.1) to 10 *Human _Heterogeneity_R5%* data and 10 *SNARE-seq Human_Heterogeneity_R10%* data (description in 3.2). Fig. 3e and f show the comparison results. Apparently, AIscEA achieved the smallest average FOSCTTM score and was more robust to its hyper-parameters. All above experiments demonstrate that AIscEA is the best method to resolve the cell-type heterogeneity problem.

### 4.2 Comparison of the cell-type alignment

In this section, we compare AIscEA with CoupleNMF [19] in terms of cell-type alignment. We applied both methods to *SNARE-seq Human, SNARE-seq Mouse 5k, SNARE-seq Mouse 10k, scMultiome-seq PBMC 3k* and *scMultiome-seq PBMC 12k*, except *SNARE-seq Human _Heterogeneity* because CoupleNMF requires the scRNA-seq and scATAC-seq data share the same number of cell types. CoupleNMF only generated results for two datasets with small number of cells, which are *SNARE-seq Human* and *scMultiome-seq PBMC 3k*. For the rest of the datasets, CoupleNMF failed and ran out of memory (memory error).

In Fig. 4, we illustrate the comparison between AIscEA and CoupleNMF on *SNARE-seq Human*. The same comparison for *scMultiome-seq PBMC 3k* is shown in Supplemantary Materials Fig. S.1(e-h). As shown in Fig. 4, for AIscEA, the cells in the aligned cell types in both scRNA-seq and scATAC-seq are well isolated. However, for CoupleNMF, the cells in the aligned cell types are mixed together. We further evaluated the performance of both methods in terms of overlapping coefficient (defined in (4)) and the Silhouette score shown in Table 2. Clearly, AIscEA achieves much higher overlapping coefficients and Silhouette scores, which demonstrates that AIscEA outperform CoupleNMF in terms of cell-type alignment.

**Fig. 4:**
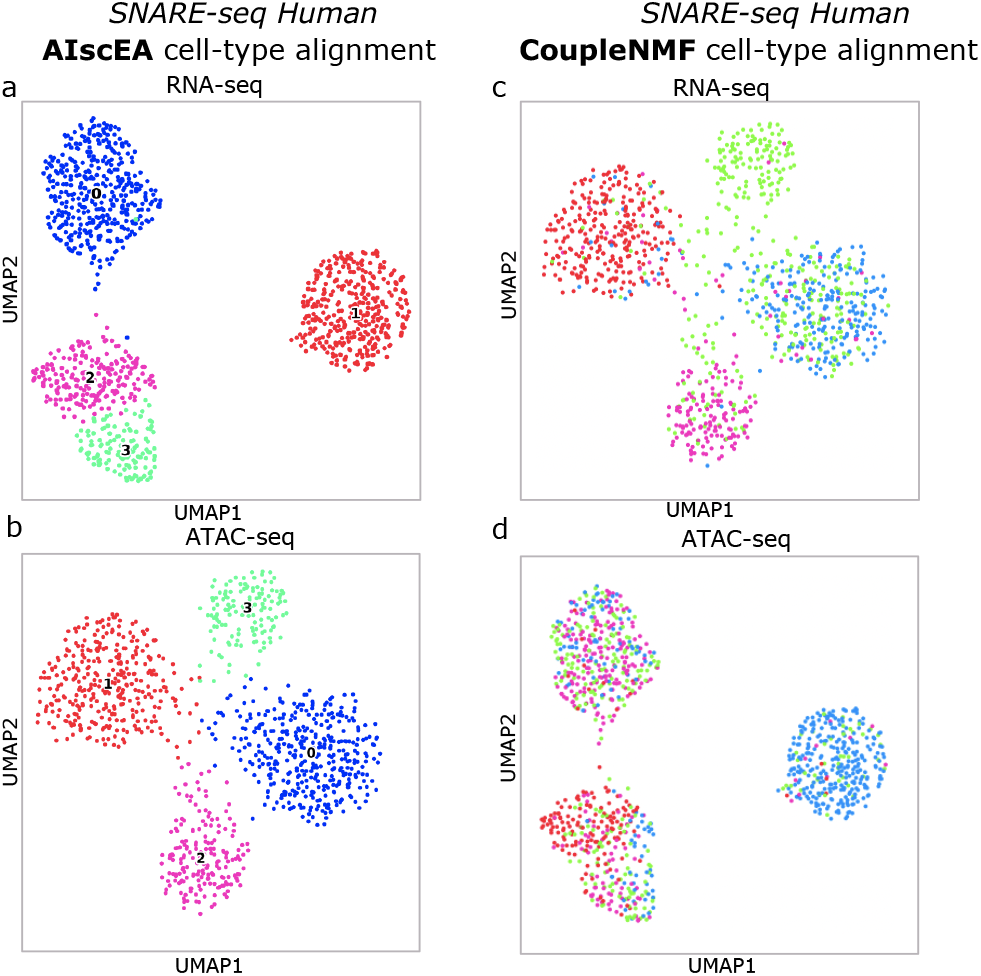
(a-b) The aligned cell types in the RNA-seq and ATAC-seq in *SNARE-seq Human* data identified by AIscEA. (c-d) The aligned cell types in the RNA-seq and ATAC-seq in *SNARE-seq Human* data identified by CoupleNMF.

**Fig. 5:**
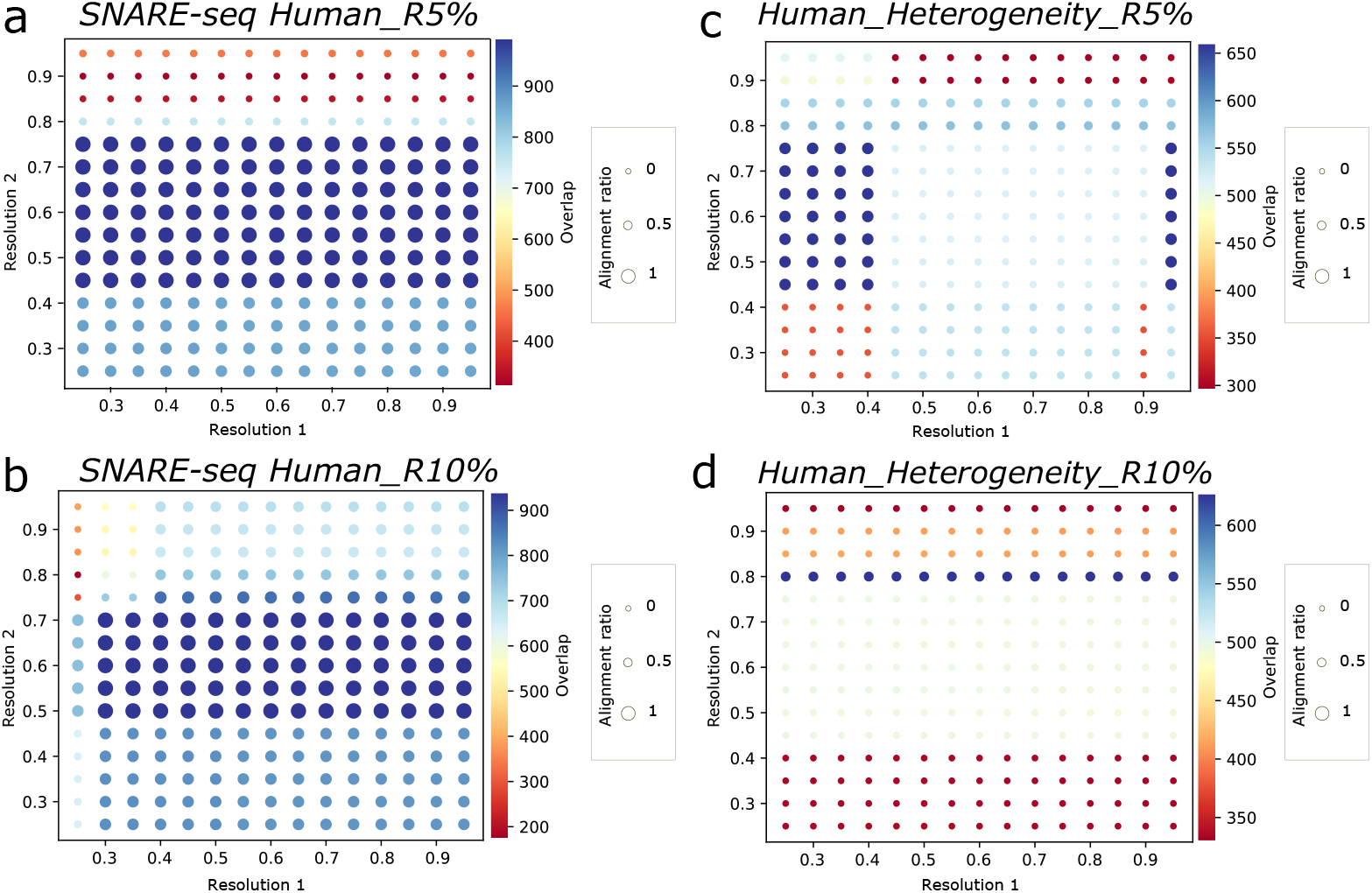
Screening the resolution parameters. Color bar denotes the number of overlapped cells identified by the cell-type alignment. Dark blue implies the cell-type alignment has good performance. The size of the dots corresponds to the *alignment ratio* defined in 2.3. Large size means the the *alignment ratio* is large. (a-b) Screening results for *SNARE-seq Human_R5%* and *SNARE-seq Human_R10%*. (c-d) Screening results for *Human_Heterogeneity_R5%* and *SNARE-seq Human_Heterogeneity_R10%*.

**Table 2:**
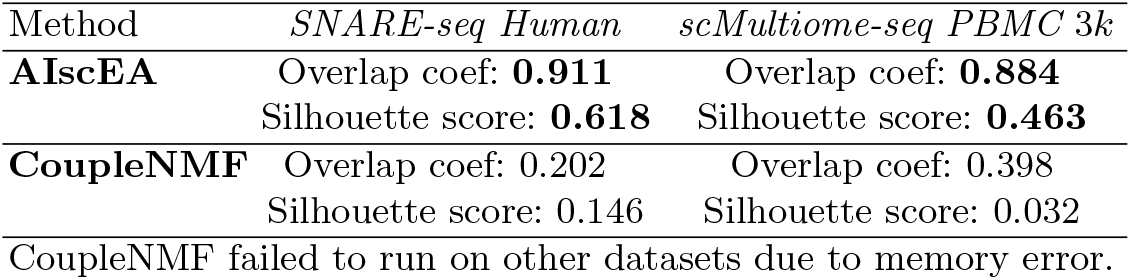
Cell-type alignment comparison.

| Method | *SNARE-seq Human* | *scMultiome-seq PBMC 3k* |
| --- | --- | --- |
| **AIscEA** | Overlap coef: **0.911**<br>Silhouette score: **0.618** | Overlap coef: **0.884**<br>Silhouette score: **0.463** |
| **CoupleNMF** | Overlap coef: 0.202<br>Silhouette score: 0.146 | Overlap coef: 0.398<br>Silhouette score: 0.032 |
| CoupleNMF failed to run on other datasets due to memory error. |  |  |

### 4.3 Comparison of cell-cell alignment using real-world data

We compared AIscEA with competing cell-cell alignment methods MMD-MA [23], UnionCom [24], SCOT [25], and Pamona [26] on both real-world SNARE-seq data and real-world scMultiome-seq data. We selected hyper-parameters for each method following the strategy we described in. Section 3.4.

#### AIscEA outperforms current methods on the real-world SNARE-seq data

We applied all competing cell-cell alignment methods on *SNARE-seq Mouse 5k* and *SNARE-seq Mouse 10k* (description in 3.2). We compared their performance in terms of the average FOSCTTM score and cell coverage (description in Section 3.3).

Fig. 6 a and b illustrate the cell-type alignment identified by AIscEA for *SNARE-seq Mouse 5k*. And Fig. 6c shows the comparison between different methods in terms of the average FOSCTTM score and cell coverage. An shown, AIscEA achieves the lowest average FOSCTTM score, which is much smaller than the rest of the methods. The cell coverage of AIscEA is slightly smaller than the other methods (4966 cells out of 5081 cells, only around 2% of cells are missed by AIscEA). But considering both average FOSCTTM score and cell coverage, it is obvious that AIscEA significantly outperforms all the current methods.

**Fig. 6:**
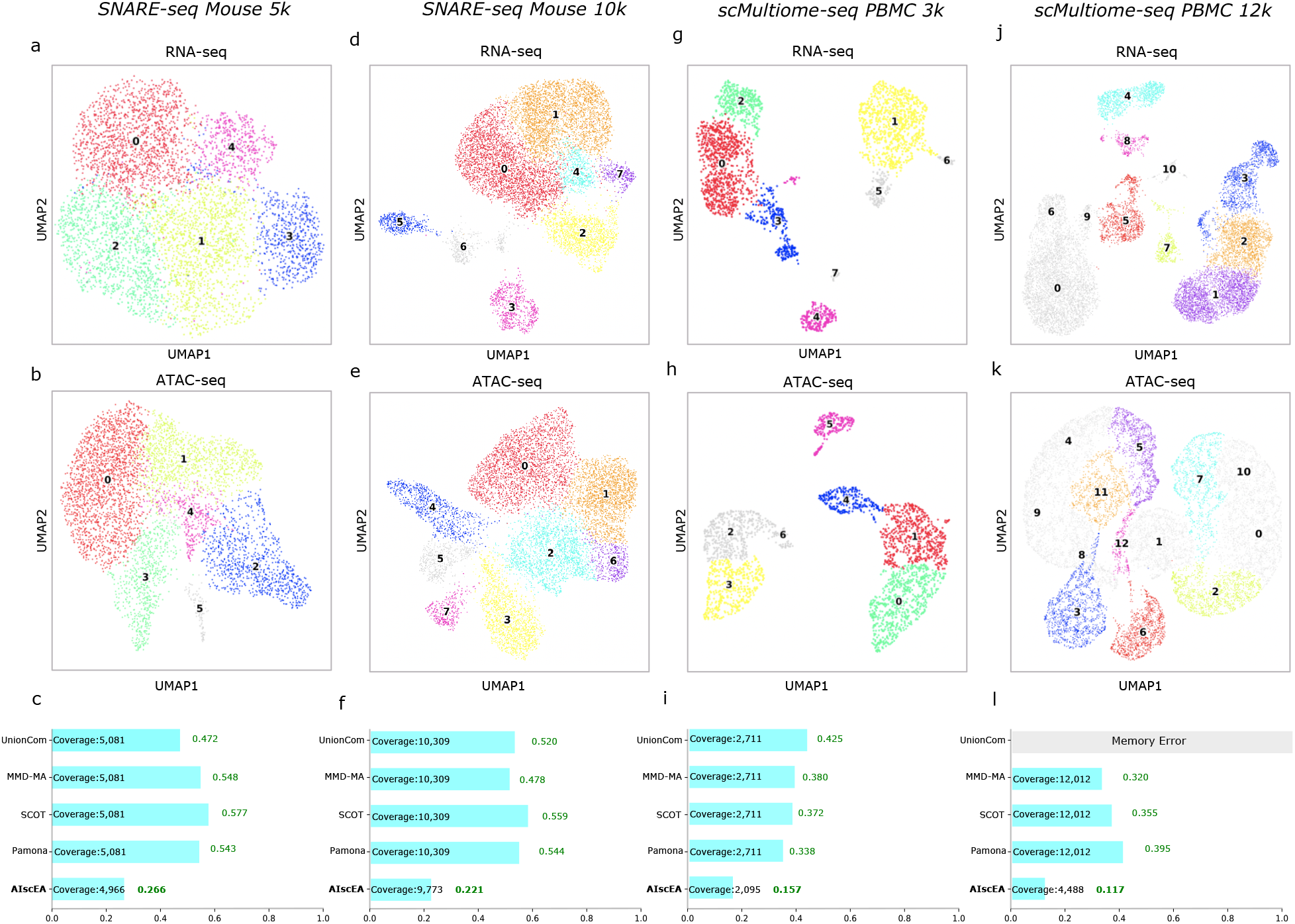
(a-b) The UMAPs of the RNA-seq and ATAC-seq of *SNARE-seq Mouse* 5*k*. Aligned cell types identified by AIscEA are shown in the same color, but grey color shows filtered out cell types by AIscEA. Each color indicates a specific aligned cell type across the measurements. (c) The bar plots of the average FOSCTTM scores for all methods. The shorter the bar the better the method performs. The number shows on the bar is the cell coverage for the method. (d-e) The UMAPs of the RNA-seq and ATAC-seq of *SNARE-seq Mouse* 10*k*. The color coding scheme is the same as (a-b). (f) The bar plots of the average FOSCTTM scores for all methods along with cell coverage for *SNARE-seq Mouse* 10*k*. (g-h) The UMAPs of the RNA-seq and ATAC-seq of *scMultiome PBMC 3k*. The color coding scheme is the same as (a-b). (i) The bar plots of the average FOSCTTM scores for all methods along with cell coverage for *scMultiome PBMC 3k*. (j-k) The UMAPs of the RNA-seq and ATAC-seq of *scMultiome PBMC 12k*. The color coding scheme is the same as (a-b). (l) The bar plots of the average FOSCTTM scores for all methods along with cell coverage for *scMultiome PBMC 12k*.

Fig. 6 d and e illustrate the cell-type alignment identified by AIscEA for *SNARE-seq Mouse 10k*. Fig. 6f shows the comparison between different methods in terms of the average FOSCTTM score and cell coverage. An shown, AIscEA achieves the lowest average FOSCTTM score, which is much smaller than the rest of the methods. The cell coverage of AIscEA is slightly smaller than the other methods (9773 cells out of 10,309 cells, only around 5% of cells are missed by AIscEA). But considering both average FOSCTTM score and cell coverage, AIscEA significantly outperforms all the current methods.

#### AIscEA outperforms the current methods on the real-world scMultiome-seq data

We applied all competing cell-cell alignment methods on *scMultiome-seq PBMC* 3*k* and *scMultiome-seq PBMC* 12*k* (description in Section 3.2). We compared their performance in terms of the average FOSCTTM score and cell coverage (description in Section 3.3).

Fig. 6 g and h illustrate the cell-type alignment identified by AIscEA for *scMultiome-seq PBMC* 3*k*. Fig. 6i shows the comparison between different methods in terms of the average FOSCTTM score and cell coverage. As illustrated, AIscEA attained the lowest average FOSCTTM score, which is much smaller than the rest of the methods. Considering both the average FOSCTTM score and cell coverage, AIscEA significantly outperforms all the current methods.

Fig. 6 j and k illustrate the cell-type alignment identified by AIscEA for *scMultiome-seq PBMC* 12*k*. Fig. 6l shows the comparison between different methods in terms of the average FOSCTTM score and cell coverage. AIscEA yielded the lowest average FOSCTTM score with a large margin. Although The cell coverage of AIscEA can be smaller than the other methods, considering both average FOSCTTM score and cell coverage, AIscEA outperforms all the current methods.

## 5 Conclusion

In this study, we proposed AIscEA, an unsupervised computational method for integrating single-cell gene expression and chromatin accessibility measurements. Unlike other approaches, AIscEA relies on the biological consistency between the two measurements to guide the integration. We compared AIscEA with the state-of-the-art methods on the *SNARE-seq* human cell line mixtures benchmark datasets [17] and demonstrated that AIscEA can effectively select hyper-parameters as well as better handle the cell-type heterogeneity problem. Furthermore, we showed that AIscEA significantly outperforms previous methods when applying to the real-world mouse SNARE-seq and scMultiome-seq datasets.

Several innovations developed in this work contributed to the performance of AIscEA. First, the ranked similarity score enables us to compare the cell types across measurements. The ranked similarity score is the key to estimating the similarity between cell types from different modalities. Second, the novel permutation test can distinguish the true cell-type alignment if the corresponding ranked similarity score is significantly larger than the random ranked similarity score in the background. Last but not least, the graph alignment method uses the symmetric *k* nearest neighbor graph to characterize the low-dimensional manifold. It is a notable advantage that AIscEA can identify cell types that appear only in one domain and exclude them from the cell-cell alignment in further analysis.

Our future direction is to recruit more cells in the integration. We believe AIscEA is the milestone for the integration of single-cell gene expression and chromatin accessibility measurements. Furthermore, it also provides a stepping stone for integrating other single-cell measurements.

