## Supplemental File for "AIscEA: Unsupervised Integration of Single-cell Gene Expression and Chromatin Accessibility via Their Biological Consistency"

### Supplemental Materials

#### A Cell-type Identification

**Preprocessing single-cell ATAC-seq** Single-cell ATAC-seq matrices are inherently larger, and they carry higher signal sparsity than RNA-seq data due to low detection efficiency [1]. Chromatin accessibility can help identify the gene regulations [2]; therefore, AIscEA detects the closest downstream genes to each accessible (open) chromatin region to estimate the gene expression activities and ATAC-seq data's potential transcriptional regulator bindings. The unique challenges in single-cell ATAC-seq downstream analysis are addressed in the preprocessing step by finding the promoter-driven regulation probability of each region in each cell. For all the experiments using AIscEA, we preprocessed the ATAC-seq data and used topic modeling framework cisTopic to get low-dimensional topics and a prediction matrix of the single-cell ATAC-seq data [3]. The prediction matrix includes the probability of each region in each cell, which is estimated by multiplying the topic-cell and the region-topic predictive distributions.

Previously, genes were classified as open if an ATAC-seq peak overlapped their promoter [4] using the BEDTools package's intersect method. AIscEA uses the BEDTools package's closest to find the open genes in the overlapping result of promoter data for the species and ATAC-seq peak regions [5–7]. Each peak is associated with an open gene using default arguments for finding the closest downstream gene considering a threshold of 1000 bp window. A region is considered distal and excluded if it lacks the closest gene within 1000 bp with promoters. Accessible Regions in cisTopic's prediction matrix were substituted with their closest downstream gene of the promoter regions to exclude low signal peaks and potential noise in the sparse ATAC-seq data. We summed up the values in the prediction matrix together for the regions that are the closest to the same gene.

**Clustering** AIscEA performs standard quality control using the single-cell RNA-seq analysis toolkits Scanpy [8] to filter out low-quality cells and genes. AIscEA normalizes the expression matrix to $10,000$ reads per cell and then takes the expression values' logarithms. In a big real-world dataset, AIscEA has an option to detect genes having high variations between cells that adopts Scanpy's *highly_variable_genes* function.

AIscEA employs Scanpy to find clusters in each domain by constructing the data's neighborhood graphs [8] on the RNA-seq data and low-dimensional topics of ATAC-seq data from cisTopic. The connectivities are estimated using Uniform Manifold Approximation and Projection (UMAP) on the $k$-nearesst neighbors graph [9] (the parameters can be tuned by the users). The graph clustering method is based on Leiden, a community detection that optimizes modularity to directly cluster the neighborhood graph of cells and provide high-quality results [8, 10]. Then, we can project the clusters found for reduced dimensional topics data, on the whole, probability prediction matrix, $\mathcal{D}$.

Similarities between clusters of different domains can be detected using co-expressed genes between pairs of RNA-seq and ATAC-seq clusters. A ranking for each cluster's differential genes is calculated by directly comparing the various clusters' gene expression distributions. AIscEA uses the differential gene expression testing, the Wilcoxon rank-sum in the Scanpy Python package [8, 11] because it is recommended as a fast nonparametric method between optimal marker selection algorithms [12] to analyze large RNA-seq data and find the marker genes [13]. For each cluster, a list of marker genes is generated along with adjusted p-values and the logarithm to basis 2 of fold change ($log2FC$) values for the marker genes. We keep each cluster's significant marker genes that their adjusted p-value is lower than 0.01 for multiple testing using the Benjamini-Hochberg correction.

First, very small clusters may be removed as they lack enough meaningful information of marker genes to form a cell type for further analyses. This step is recommended when the users aim to align cells in an extensive real-world dataset containing much noise.

### B Cell-cell Alignment via Graph Alignment

#### B.1 Construction of the Symmetric $k$ Nearest Neighbor Graph

Based on [14], we know that the $k$ nearest neighbor graph might not be able to present low-dimensional geometry. Specifically, it prefers to go through regions of low density and even takes large detours in order to avoid the high density regions. To overcome the problem, we propose the symmetric $k$ nearest neighbor graph.

Let $A$ be the adjacency matrix a of $k$ nearest neighbors graph $\mathcal{G} = (V, E)$. We construct a new graph whose adjacency matrix is $B = I(A + A^2)$, where $I$ is a element-wise binarize function where $I(x) = 1$, if $x > 0$, and otherwise $I(x) = 0$. Then we keep the edges between nodes $i$ and $j$ in the new graph, if and only if $B_{ij} = B_{ji} = 1$. We call the new graph symmetric $k$ nearest neighbor graph.

#### B.2 The Frank-wolf Algorithm

To solve the following optimization, we proposed to use Frank-Wolfe algorithm to solve it.

$$\begin{aligned} &\max_{P} : \operatorname{Tr}(A_1^\top P A_2 P^\top) + \lambda \operatorname{Tr}(PL) \\ &s.t.\ P \in \{P \in \{0,1\}^{N\times N}, P^\top \mathbf{1}_N = \mathbf{1}_N, P\mathbf{1}_N = \mathbf{1}_N\}. \end{aligned} \tag{S1}$$

The Frank-Wolfe algorithm is as follow.

**Algorithm 1**: Frank-Wolfe using Line Search for the step size $\gamma$

Let $\boldsymbol{P}^{(0)} \in \mathcal{P}$ ▷ $\boldsymbol{P}^{(0)}$ is randomly selected from set $\mathcal{P}$ ;
**while** $k \leq K$ **do**
  $\boldsymbol{s} \leftarrow \arg\min_{\boldsymbol{s}\in\mathcal{D}} \left\langle \boldsymbol{s}, \nabla f\left(\boldsymbol{P}^{(k)}\right)\right\rangle$;
  $\gamma \leftarrow \arg\min_{\gamma\in[0,1]} f\left(\boldsymbol{P}^{(k)} + \gamma\left(\boldsymbol{s} - \boldsymbol{P}^{(k)}\right)\right)$ ;
  $\boldsymbol{P}^{(k+1)} \leftarrow (1-\gamma)\boldsymbol{P}^{(k)} + \gamma \boldsymbol{s}$ ;
**end**
**return** $\boldsymbol{P}^{(k+1)}$ ▷ Optimal permutation matrix ;

In the algorithm, $\nabla f\left(\boldsymbol{P}^{(k)}\right) = \lambda A_{\mathcal{G}_1}^T P A_{\mathcal{G}_2} + (1-\lambda)L^T$. The optimal step size $\gamma$ can be computed as follow.

$$\gamma^* = -\frac{1}{2}\frac{\operatorname{tr}(T_2)}{\operatorname{tr}(T_1)}, \tag{S2}$$

where

$$\begin{aligned} T_1 &= \frac{\lambda}{2} A_{\mathcal{G}_1}^T \left(s - x^{(k)}\right) A_{\mathcal{G}_2} \left(s^\top - \left(x^{(k)}\right)^T\right), \\ T_2 &= (1-\lambda)\left(S - x^{(k)}\right) L + \frac{\lambda}{2}\left[x^{(k)} A_{\mathcal{G}_2}\left(S^T - \left(x^{(k)}\right)^T\right) + \left(s - x^{(k)}\right) A_{\mathcal{G}_2}\left(x^{(k)}\right)^\top\right] \end{aligned} \tag{S3}$$

#### B.3 The Path-relinking Algorithm

Since the objective function of (S1) is not convex, the Frank-Wolf algorithm cannot find the optimal solution. In order to find a better solution, we propose to use the path-relinking strategy. First, we use the Frank-Wolf algorithm to quickly find several local optimal solutions. Then we try to link these solutions with the hope that on the linking path, better solution can be identified. The path-relinking strategy has been used in [15].

### C Experimental Settings

#### C.1 Datasets

**SNARE-seq Human** SNARE-seq data [16] is generated by a robust droplet-based method that simultaneously measured the transcriptome and accessible chromatin in single cells. Mixtures of cultured human BJ, H1, K562, and GM12878 cell types are collected to generate 1047 paired profiles in Gene Expression Omnibus accession GSE126074. They captured both single-cell chromatin information and mRNA expression values of the same cell populations. Gene expression data is provided as a count matrix of size $1047 \times 18666$ where rows of the matrix are single cells, and the columns are the captured genes. The entries of the matrix are RNA-seq captured values for each gene in each cell. Chromatin information can be found in a single-cell ATAC-seq binary matrix of size $1047 \times 136771$. Similar to gene expression data, the rows represent cell tags in ATAC-seq data. Although, the columns of the ATAC-seq matrix show accessible chromatin regions (peaks) for the cells.

To further test the robustness of each method in terms of hyper-parameters, we generate 10 datasets from the SNARE-seq Human dataset, where 5%, 10% are randomly removed.

**SNARE-seq Human_heterogeneity** We experimented with removing a cluster from one domain in SNARE-seq Human data to analyze AIscEA's performance when ATAC-seq has a domain-specific cluster that is not shared in RNA-seq data. The size of RNA-seq data after removing one cluster will be $711 \times 18666$, which form three clusters, and the ATAC-seq data stays the same as the original SNARE-seq Human data, a binary matrix of size $1047 \times 136771$. We named this dataset *Human_heterogeneity* as it simulates heterogeneous cell types between two domains.

To further test the robustness of each method in terms of hyper-parameters for datasets with cell-type heterogeneity problem, we generate 10 datasets from the SNARE-seq Human_Heterogeneity dataset, where 5%, 10% are randomly removed.

**Two real-world SNARE-seq datasets** SNARE-seq of adult mouse cerebral cortices provides dual-omics profiling of $10,309$ paired profiles from adult mouse cerebral cortices. They captured 12 replicates and 22 cell clusters, including ten excitatory neuron types, four inhibitory neuron types and OPCs, newly formed Itpr2, and mature oligodendrocytes, as well as other non-neuronal cells [16].

Moreover, SNARE-seq of neonatal mouse cerebral cortex captured 5 replicates of dual-omics data in neonatal mouse cerebral cortex. The profiling included 5,081 cells that linked transcriptome and chromatin accessibility regions

**Two 10x PBMC datasets** We used data from a healthy donor - granulocytes removed through cell sorting (3k) that includes Cryopreserved human peripheral blood mononuclear cells (PBMCs) [17]. According to Single Cell Multiome ATAC + Gene Expression Sequencing, Granulocytes were removed by cell sorting. Isolated nuclei were captured based on Chromium Next GEM Single Cell Multiome ATAC + Gene Expression user guide (CG000338 Rev A) and sequenced on Illumina Novaseq 6000 v1 Kit. This dataset includes an estimated number of 2711 cells and can be found on the 10x website. They have detected $25,805$ total genes and $98,319$ peak region in single-cell RNA-seq and ATAC-seq, respectively.

We also employed a larger PBMC dataset from a healthy donor - granulocytes removed through cell sorting (12k) from 10x website [18]. This dataset consists of $29,717$ genes in RNA-seq and $143,887$ peak regions in ATAC-seq dataset for $11,898$ single cells.

#### C.2 Hyper-parameter selection

We used the range of hyper-parameters suggested by the authors of each method to find the optimal set that yields the best FOSCTTM score for SNARE-seq Human dataset as well as Human_heterogeneity dataset.

**MMD-MA** For the MMD-MA method [19, 20], the grid search consists of three or four hyper-parameters to be tuned based on two pipelines. The more recent MMD-MA pipeline has three hyper-parameters as one hyper-parameter $b$ can be estimated using their pipeline. The Gaussian's width for the initial kernel calculation over each data point $\sigma \in \{0.01, 0.1, 1.0, 10\}$ is an optional hyper-parameter because it can be estimated in more recent MMD-MA method. The bandwidth, $\sigma$, effectively controls the dimensionality of the Hilbert space induced by the kernel. Relative weights $\lambda_1, \lambda_2 \in \{10^{-3}, 10^{-4}, 10^{-5}, 10^{-6}, 10^{-7}\}$ for the terms in the optimization terms are two other hyper-parameters. The fourth hyper-parameter is the dimensionality of the latent space $p \in \{3, 4, 5\}$.

**UnionCom** In the UnionCom [21] pipeline, we need to specify four hyper-parameters including: the number of neighbors in the graph $k \in \{5, 10, 25, 50, 75, 100, 150, 200, 250, 300\}$, the embedded space dimensionality $p \in \{2, 5, 10\}$, the trade-off for the embedding $\beta \in \{0.001, 0.005, 0.01, 0.5, 0.1, 0.5, 1, 5, 10\}$, and finally a regularization coefficient $\rho \in \{0.001, 0.005, 0.01, 0.5, 0.1, 0.5, 1, 5, 10\}$.

**SCOT** SCOT [22] requires two hyper-parameters to be tuned: the regularization weight $\epsilon \in \{0.0005, 0.001, 0.01, 0.1, 1, 10\}$ and the number of neighbors $k \in \{10, 20, 30, 40, 50, 75, 100, 125, 150, 175\}$.

**Pamona** Pamona [23] requires four hyper-parameters to be explored in the grid search. The number of neighbors in the $k$-nearest neighbors graph $k \in \{10, 20, 30, 50, 75, 100\}$, the regularization parameter of the partial-GW framework $\epsilon \in \{0.0001, 0.001, 0.01, 0.1\}$, the parameter $\lambda \in \{0.5, 0.75, 1.0, 1.5\}$ of manifold alignment for the trade-off between aligning identical cells and preserving the local topology, and output dimension of the shared embedding space after the manifold alignment $p \in \{10, 20, 30, 40, 50\}$. For the Pamona method, we explored two different scenarios. First, we ran the original pipeline. Second, we provided the actual number of shared cells as prior information as they claimed this would improve the method's performance. In the analysis, Pamona(v2) refers to the case when providing the true number of shared cells between two domains as prior information provided for the method.

##### C.3 CoupleNMF

We compared AIscEA cell-type alignment with CoupleNMF [24], a state-of-the-art method for cell-type alignment across domains. The coupleNMF method is not fully unsupervised as it requires the number of cell types in the data as an input, which is unknown in most problems. In addition, it consumes high memory. Because of these two reasons, we could only run it on two smaller datasets that the true number of cell types has been recommended. We then ran CoupleNMF method using the prior knowledge of the number of clusters. We kept the rest of hyper-parameters as default, as CoupleNMF has not provided an automatic hyper-parameter tuning. AIscEA showed superior performance and more well-defined clusters as shown Fig. S1. Furthermore, Table 2 shows higher overlap coefficient and silhouette score in AIscEA, in comparison to CoupleNMF. AIscEA consistently yielded better scores in cell-type alignment and showed high robustness.

##### C.4 AIscEA

We explored hyper-parameters for cell-type alignment as shown in Fig.S2: resolution values in the range of $\{0.1, 0.15, 0.20, ..., 1.5\}$.

As we discussed in the main paper, after cell-type alignment, we also screened hyper-parameters for cell-cell alignment: 1) $\lambda \in \{0.1, 0.2, 0.3, 0.4, 0.5, 0.6, 0.7, 0.8, 0.9\}$, and 2) $k = \{3, 6, ..., 100\}$.

##### C.5 AIscEA selects hyper-parameters that preserve biologically meaningful mapped clusters in cell-type alignment.

We screened all hyper-parameters for AIscEA in the cell-type alignment procedure to analyze the effectiveness of our heuristic hyper-parameter selection. This step addresses the robustness of the initial clustering and its impact on the final result. The exact knowledge of clusters depends on the biological question of interest, which is inaccessible in most real-world data. Still, if the true knowledge is not provided, we extract the prominent clustering of each domain and estimate the primary clusters. Each primary cluster may consist of one or some biologically similar cell types. Fig. S2(a-d) shows the screening of $resolution1$ and $resolution2$ for downsampled SNARE-seq Human and SNARE-seq Human Heterogeneity. Our hyper-parameters $resolution1$ and $resolution2$ automatically determine the clusters. The size of each circle in the dot plots shows the heuristic values "mapped cluster ratio" for one single experiment. We observed that the bigger the circles are, the darker blue they appear. Therefore, the unsupervised feature "alignment ratio" serves as an estimator for the overlapping number of cells between two domains when the data lacks cell barcodes. We showed a subset of grid search values experimented for ease of visualization. Similar dot plots for real world datasets of *SNARE-seq Mouse* $5k$ *and* $10k$ as well as *PBMC* $3k$ *and* $10k$ are also shown in Supplementary Materials.

##### C.6 Optimal hyper-parameters for SNARE-seq Human and Human_Heterogeneity dataset

We report the optimal hyper-parameters for SNARE-seq Human dataset as well as Human_Heterogeneity dataset.

**Table S1.** Best parameters for SNARE-seq Human dataset as well as Human_heterogeneity dataset after screening all the hyper-parameters. For AIscEA, $resolution1$ and $resolution1$ are shown as $r1$ and $r2$ for ease of visualization.

| Method | Hyper-parameters Human | Hyper-parameters Human_heterogeneity |
|---|---|---|
| SCOT | $k = 50$, $e = 0.0005$ | $k = 125$, $e = 10^{-3}$ |
| MMD-MA | $l1 = 10^{-7}$, $l2 = 10^{-6}$, $p = 5$, $(b = 0.1)$ | $l1 = 10^{-3}$, $l2 = 10^{-7}$, $p = 3$, $(b = 1)$ |
| Pamona | $e = 10^{-3}$, $k = 75$, $l = 0.5$, $dim = 10$ | $e = 10^{-3}$, $k = 10$, $l = 0.5$, $dim = 2$ |
| UnionCom | $n = 5$, $dim = 2$, $\sigma = 10^{-3}$, $r = 0.5$ | $n = 25$, $dim = 5$, $\sigma = 0.01$, $r = 5$ |
| AIscEA | $\lambda = 0.8$, $k = 3$, $r1 = 0.25$, $r2 = 0.45$ | $\lambda = 0.8$, $k = 3$, $r1 = 0.25$ , $r2 = 0.45$ |

#### D Method Evaluation

We compare AIscEA with four state-of-the-art single-cell alignment methods, Pamona, SCOT, MMD-MA, and UnionCom. None of the methods utilize any correspondence information either between cells or between features during the alignment. However, Pamona, SCOT, MMD-MA, and UnionCom require cell correspondence information for selecting the best set of hyper-parameters for cell alignment.

We ran all methods over a hyper-parameters tuning grid search and chose the setting that generated the optimal score. The metric to measure the score is called the "fraction of samples closer than the true match" (FOSCTTM), which has been used to measure the quality of the cell alignments [21]. AIscEA does not use the correspondence information between cells in the alignment process, and it only uses them to report the performance after the alignment is generated.

##### D.1 FOSCTTM

In a perfect cell alignment, each cell in the first domain is aligned to another cell in the second domain with the least distance to its correct match in the second domain. We compute the Euclidean distance between the corresponding cell (in the second domain) of each cell (in the first domain) to every other cell in the second domain. After ranking the distance values, we can find the fraction of distances closer to that cell than its true match. FOSCTTM score is calculated by the average of the fraction values for all the cells. An excellent alignment method must yield a zero value of FOSCTTM. The lower the FOSCTTM score is, the more reliable the method performs.

MMD-MA and UnionCom project both datasets to a shared latent space to calculate the FOSCTTM score, but SCOT and Pamona project one dataset onto the other. We project RNA-seq cells on ATAC-seq space and calculate FOSCTTM scores accordingly. AIscEA calculates the FOSCTTM score by projecting

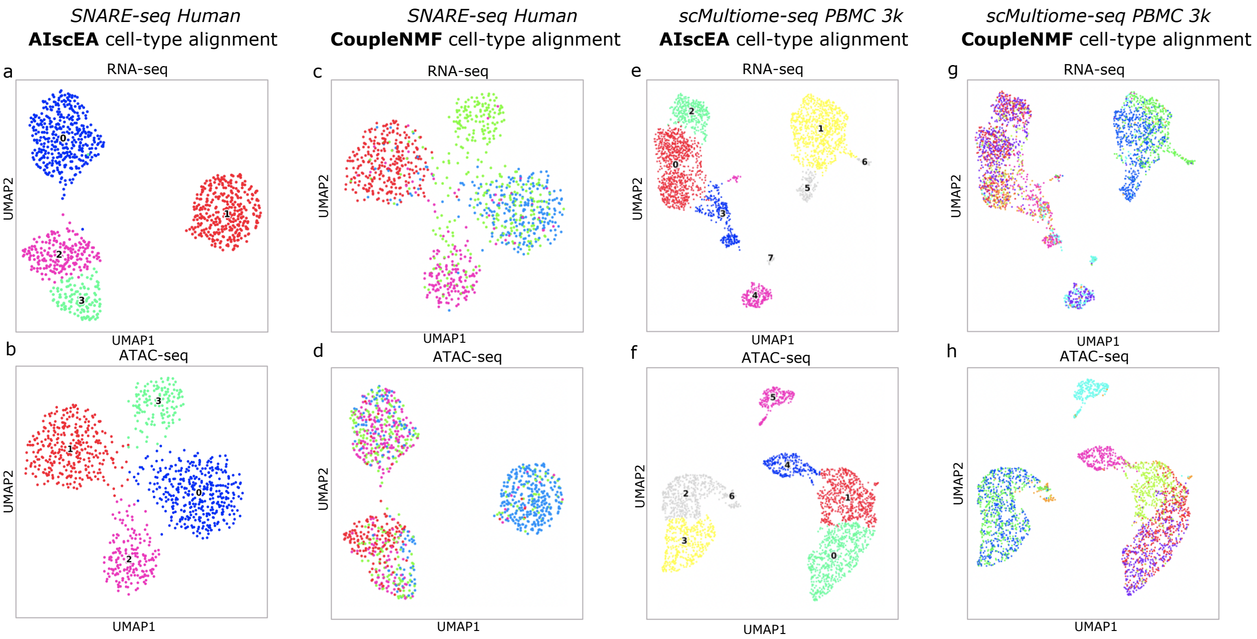


**Fig. S1.** (a-b) The aligned cell types in the RNA-seq and ATAC-seq in *SNARE-seq Human* data identified by AIscEA. (c-d) The aligned cell types in the RNA-seq and ATAC-seq in *SNARE-seq Human* data identified by CoupleNMF. (e-f) The aligned cell types in the RNA-seq and ATAC-seq in *PBMC 3k* data identified by AIscEA. (e-f) The aligned cell types in the RNA-seq and ATAC-seq in *PBMC 3k* data identified by CoupleNMF. The aligned cell types are in the same colors across the RNA-seq and ATAC-seq.

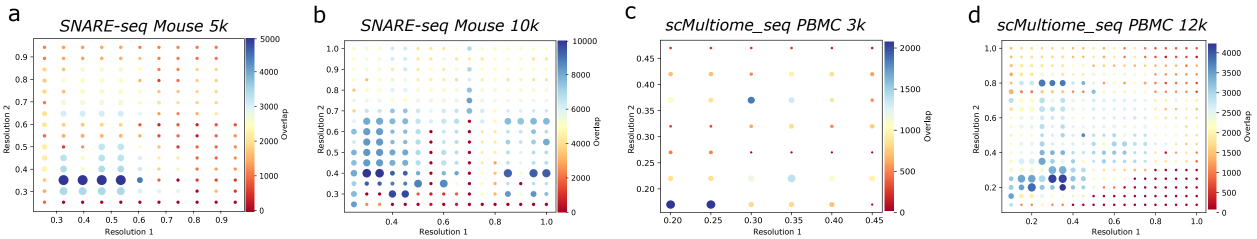


**Fig. S2.** (a) Grid-search over hyper-parameters used in cell-type alignment. Correlation between number of overlapping cells and "alignment ratio" in RNA-seq and ATAC-seq data in grid search over resolution1 and resolution2 to get the initial clusters in (a) *SNARE-seq Mouse 5k*, (b) *SNARE-seq Mouse 10k*, (c) *scMultiome-seq PBMC 3k*, and (d) *scMultiome-seq PBMC 12k*. Size of the dots is correlated with the score "mapped cluster ratio" for each pair of resolution 1 and resolution 2. Maximum number of possible overlapping cells differs in each of four experiments, the highest number is indicated in dark blue in the color bar. We ran a higher range of hyper-parameters, but only a subset of them are shown in this figure for ease of visualization.

cells from one domain to their exact mapped cells in the second domain to find the fraction of cells closer than the true match. Therefore, for all cells in scRNA-seq data, we first project them to their mapped cells in scATAC-seq data. Then, we find how many cells are closer to the mapped cells in scATAC-seq data than the true match in scATAC-seq data. The true match in scATAC-seq dataset has the same cell barcode as the cell of interest in scRNA-seq dataset. This idea can be generalized to use super-cells in our method by calculating the fraction of cells for all the cells in the super-cell.

#### D.2 Overlap coefficient

Furthermore, we used the overlap coefficient to show the overlapping cells between the clusters that are mapped together in the embedded space after the alignment process. CoupleNMF clusters are used with the embedded UMAP values. Then, cell barcodes overlap and union for each cluster can be calculated to find overlap coefficients.
